# Enhancing cassava reproductive development does not negatively impact shoot to root ratio and dry matter content in storage roots

**DOI:** 10.1101/2020.12.19.423574

**Authors:** Oluwasanya Deborah, Setter Tim

**Author notes:** **Correspondence:** Tim Setter.

## Abstract

From previous studies we developed treatments that significantly improved cassava female flower and fruit development, using a combination of the anti-ethylene, silver thiosulfate (STS), and the cytokinin, benzyladenine (BA); collectively referred to as plant growth regulators (PGR). In this study, we investigated whether the benefit derived from this treatment altered partitioning of photosynthate to other sinks and general vegetative growth of cassava in the first six months of plants growth, when reproductive growth initiates and peaks. Our flower enhancing treatment did not significantly alter shoot and storage root fresh weight, partitioning index on a fresh weight basis and percent dry matter content of storage roots between Months 2 and 5. With the onset of the dry season in Month 6, PGR treated plants had higher shoot and storage root fresh weight than controls but these plant parts responded proportionally and so partitioning index between controls and treated plants was not significantly different. The nighttime starch export under PGR treatments was reduced at Months 2, 4 and 5 but this was not correlated with flower development at these months. The survival of PGR treated plants until harvest was however reduced owing to increased mortality arising from phytotoxicity and increased susceptibility to disease. We therefore conclude that PGRs have effects more directly on flower and fruit reproductive signaling and regulatory pathways rather than on an indirect effect on resource partitioning.

## 1 Introduction

Cassava (*Manihot esculenta*) is a tropical perennial plant grown as an annual for its edible starchy roots (Liu et al. 2014). It is an important staple food in tropical countries and one of the top five most important sources of starch globally (Jansson et al. 2009). Although it is asexually propagated for crop production, there is also a need for sexual reproduction in cassava crop breeding and improvement via the application of genomic selection, marker assisted breeding, and other contemporary methods (Wolfe et al. 2017; Ceballos et al. 2015). Sexual reproduction is therefore necessary for genetic recombination.

While there is opportunity to use newly-developed breeding and genetic techniques to increase the rate of genetic improvement of cassava, such efforts are constrained by cassava’s late flowering and poor flower and seed production (Ceballos et al. 2020). Poor flowering slows breeding progress and it limits the ability to use certain poor-flowering or non-flowering lines as parents in crosses (Ceballos et al. 2020). Research and development led by the Nextgen Cassava program (https://www.nextgencassava.org/), to which the present research is associated, has identified techniques to improve flower induction and flower/seed productivity. These techniques include extended daylength using red LED lights powered by solar panels to hasten the time of flowering (Pineda et al., 2018), and application of plant growth regulators (PGR) to increase flower production (Abah et al., 2016; Abubakar et al., 2016). To improve cassava flower production, the anti-ethylene, silver thiosulfate (STS), and the cytokinin, benzyl adenine (BA), are promising (Abah et al., 2016; Hyde et al., 2020, Chapter 3 of this thesis). Current research is testing various methods and timings of application and dosages. However, further technology development for PGR treatments that improve flower production would benefit from a better understanding of their mechanisms of action in cassava. In addition to direct effects on molecular signaling and regulatory systems for flower development, it is possible that STS and BA indirectly affect flowering by modifying the activities of other processes.

Cassava is a storage-root crop that stores a large proportion of its photosynthate in the harvested storage root. The harvest index of dry matter in cassava is typically about 60% or more (Munyahali et al. 2017). Cassava initiates flowering at the same or later time frame as it initiates storage root production. Cassava continues active leaf and stem growth after storage root growth is initiated, and roots grow steadily in diameter and length for an extended period of time, storing starch in each new region of root cell proliferation and expansion (El-Sharkawy and Cock 1987; Connor et al. 1981; Mehdi et al. 2019; Fernie et al. 2020). Simultaneously, plant height and new leaves are produced such that both below-ground and above-ground sink activity occur at the same time (El-Sharkawy and Cock 1987). Under this circumstance, it is possible that the poor flowering in cassava could be due to an inability of reproductive organs to compete for photosynthate. Furthermore, the perennial nature of cassava is possibly reflected in the late and poor flowering nature of genotypes that tend to have agronomically valuable storage root traits.

We have previously shown that the combination of PGRs silver thiosulfate and benzyladenine significantly improved female flower and fruit development (Chapter 3). In this study, we considered whether PGR treatments provide benefit to flowers and fruits by altering photosynthate partitioning among the other sinks – the root and shoot vegetative parts of cassava. We followed the reproductive development, whole plant growth, and dry matter content in storage roots and carbohydrate exported from leaves for the first six months after planting under PGR treatment relative to an untreated control. Our findings indicate that PGR treatment did not significantly change plant growth generally but reduced nighttime starch export in specific months. This reduction in nighttime export while storage root and vegetative growth was unaltered was, however, contrary to the hypothesis that PGR treatments limit competition from roots and vegetative plant parts and was responsible for the observed improvement in reproductive growth.

## 2 Materials and Methods

### 2.1 Field conditions

All experiments were conducted under field conditions at the International Institute of Tropical Agriculture (IITA), Oyo State, Ibadan (7.4° N and 3.9°E, 230m asl). The soil was an Alfisol (oxicpaleustalf) (Moormann et al. 1975). The land was tilled and ridged with 1 m spacing; plants were sown on top of the ridge. The field in 2018 was previously planted with maize (*Zea mays*) while the field in 2019 was previously planted with yam (*Dioscorea rotundata*) but allowed to fallow for a year before the experiment was planted; no extra nutrients or soil amendments were added to the soil. Fields were kept free of weeds with manual weeding.

### 2.2 Plant materials and treatments

Three genotypes, representing three flowering times and extents of flower profuseness were used for plant growth regulator (PGR) studies. These were IITA-TMS-IBA980002 (early and profuse), IITA-TMS-IBA30572 (middle genotype) and TMEB419 (late and poor flowering). Storage root development was examined by monthly destructive harvest under PGR and control treatments. The main experiment was conducted between June and December of 2019.

Examination of the effect of PGRs on cassava survival and storage at harvest was conducted using a subset of data on an experiment previously conducted to study the effect of PGR on cassava reproductive development (Oluwasanya et al.2020). This experiment was conducted in two crop cycles between June and December of 2018 and 2019.

### 2.3 Plant growth regulators and method of application

Silver thiosulfate (STS) was prepared by mixing 1 part 0.1M silver nitrate (AgNO_3_) dropwise with 4 parts 0.1M sodium thiosulphate (Na_2_S_2_O_3_), yielding a 20 mM stock solution. The stock solution was diluted with distilled water to 2mM. Benzyladenine (BA) solution was prepared by diluting a 1.9% (w/v) BA stock (MaxCel®, Valent BioSciences Corporation, Libertyville, IL, USA) with distilled water to 0.5mM. BA was applied by spray (about 5 mL) to the shoot apex, every seven days while a mixture of equal volumes of STS and BA was applied by “petiole feeding” every 14 days. In the petiole feeding method, the leaf blade was removed using a surgical scissors and the petiole was inserted into a 15-mL conical-bottom centrifuge tube (Falcon Brand, Corning, NY, USA) containing 10 mL of PGR solution. PGR was taken up via the petiole into xylem from which it was distributed internally to target organs in the leaves and apex. Petioles were allowed to remain immersed in PGR solution for 72h after which tubes were removed. On weeks with petiole feeding, spray treatments were applied 24 h after petiole treatments. PGR treatments were initiated six weeks after planting.

### 2.3 Data collection

To examine the monthly effect of PGR on reproductive and storage root development, the experiment comprised of five plots. Each plot was split by genotype and each genotype subplot was split by treatments (PGR vs Control). In addition, each treatment sub-subplot was comprised of a row dedicated to scoring reproductive development with borders and two rows for destructive monthly sampling to obtain storage root data as above. The maximum fruit count for each plant over four-week periods was analyzed as the representative of reproductive response to treatment per time. Data for reproductive and storage root development were collected over a 24-week period. All data was recorded using Field Book software application (Rife and Poland 2014).

We also measured starch and total sugars accumulated in leaves in the evening (1600hrs) and the amount that remained in the morning (800hrs) after exports at night. On a monthly schedule, discs (5 mm diameter, three per leaf) from the fully expanded leaf nearest to the highest shoot apex were sampled into 80% methanol and stored at −20°C until all samples were collected.

Sampling began at 8 weeks (2 months) after planting (24hrs before harvesting), until the 24th week after planting. One plant per genotype and treatment was sampled from four plots each month. Carbohydrate extraction and quantification proceeded as described by Setter et al. (2001) with modifications described by Setter and Parra (2010).

Harvest data of plants at about 12 months after planting was obtained from 6 blocks arranged in a randomized block design, each containing 8 plants per genotype per treatment from the experiment conducted in 2018 and 2019.

### 2.4 Statistical Analysis

Storage root traits and carbohydrate measurements were modeled as a linear mixed model with treatments as the independent variable while count data such as the numbers of fruits and storage roots were modelled using the negative binomial model. Flowering time was subjected to survival analysis using the Kaplan-Meier Model (Bland and Altman 1998). Due to similarities of genotypic response to PGR the means across all genotypes are reported here. Individual genotypic responses are reported in the supplementary data.

Models were built using the “Tests in linear mixed effects models” (lmerTest) (Kuznetsova et al. 2017) and the Generalized Linear Mixed Models using Template Model Builder (glmmTMB) (Brooks et al. 2017) packages in R (Team 2013). Sources of variation for reproductive growth and monthly harvest were treatment and month of sampling, while for 12 month harvest it was treatment and plant status (whether it survived and or had harvestable roots). In both cases plot was modeled as a random effect. The emmeans package (Lenth 2019) was used for post-hoc tests. Multiple means comparison was carried out using the Tukey-HSD method.

## 3 Results

### 3.1 Cassava reproductive development over time

PGR treatments did not significantly affect flowering time or the likelihood of flowering (Figure 1a). There were, however, significantly fewer males and correspondingly, more females (Figure 1b-d). The PGR treatment increased the total number of flowers per plant and increased the proportion of flowers that were female from about 19% in the control to 91% in the PGR treatment (average for the period from Month 4-5). The larger number of female flowers in the PGR treatment was associated with more fruits in PGR treated plants than in the control (Figure 1b-d).

**Figure 1.**
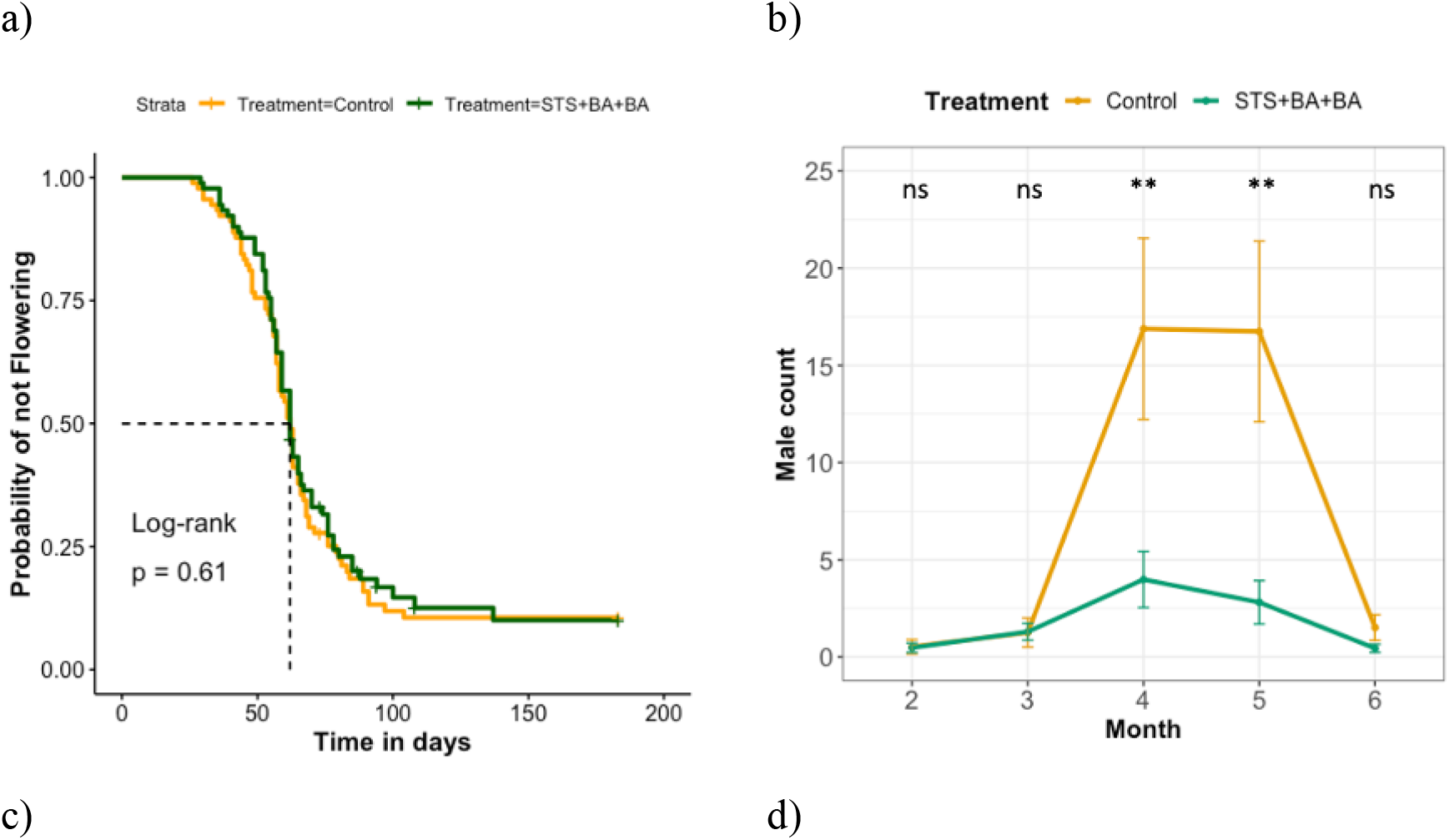
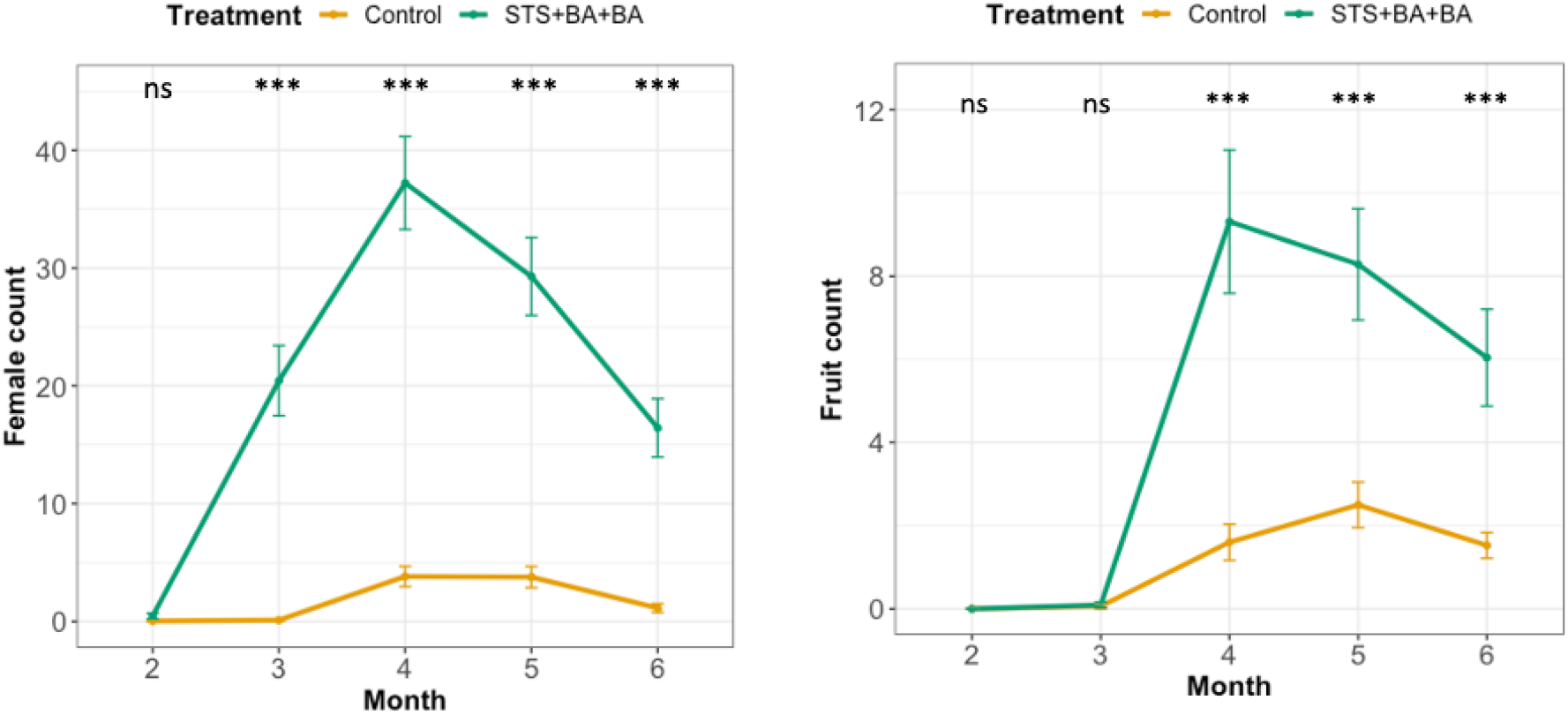
Cassava reproductive development over time in PGR treated vs Control. a) Likelihood of flowering, Kaplan-Meier curves for flowering times. b) Maximum male flowers c) Maximum female flowers. d) Maximum fruit counted per month. *, ** and *** indicate statistical significance on pairwise comparisons between treatments at 0.05, 0.01, and 0.001 significance levels, respectively.

### 3.2 Cassava vegetative growth over time

Shoot and storage root fresh weights increased throughout study period but was not significantly different between control and PGR treatment except in Month 6 (Figure 2 b, c). In Month 6, shoot weight and storage root fresh weight were significantly higher with PGR treatment than the controls. This increase in shoot and root growth of PGR plots was associated with more lateral branches on PGR treated plants that generally had a bushy appearance as a result of cytokinin contained in treatment which activates repressed buds. Partitioning index (PI, calculated as the ratio of storage root to total plant weight on a fresh weight basis) and storage root percent dry matter content (DMC) increased from Month 2 to 4, then plateaued at Month 5 and 6 (Figure 2 c, d). Initiation of storage root growth was apparently initiated between Month 2 and 3, and as storage roots accumulated starch from Month 2 to 5, the water content decreased, and percent DMC increased.

**Figure 2.**
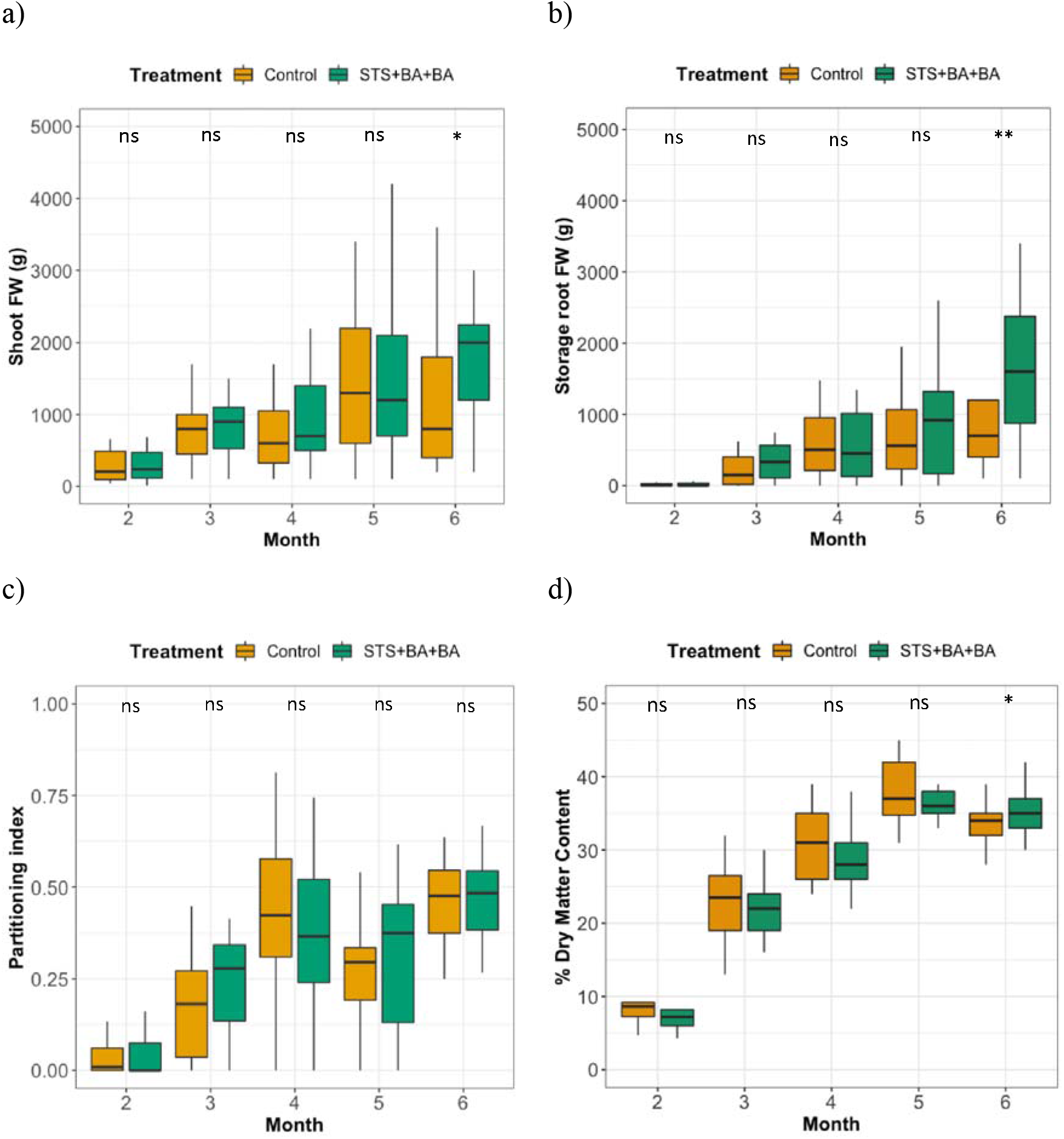
Vegetative growth over time over time in PGR treated vs Control. a) Shoot fresh weight b) Storage root fresh weight c) Partitioning index (on a fresh weight basis, see text) d) Percent dry matter content. *, and ns indicate statistical significance or no statistical significance, respectively, on pairwise comparisons between treatments (P<0.05).

### 3.3 Carbohydrate accumulation and export over time

The total sugars and starch contents per unit leaf area in sampled recently matured leaves was determined at two daily time points to provide an indication of source-sink status based on the extent of photosynthate accumulation during the day, and export at night. Other contributors to the nighttime decline include respiratory losses and metabolism to other compounds. For sugars, the general pattern was that early in the season (Month 2), leaves accumulated substantial sugar during the day, then as storage root growth ensued (Month 3 to 6), the extent to which leaves accumulated sugar was much less (Figure 3a). In the morning, sugar content had become depleted to low basal levels of about 75 μg/cm^2^; this level was consistent at all stages of crop growth (Figure 3b). In general, PGR treatment did not affect total sugar content in leaves at 16:00 pm (evening), which reflects accumulation from photosynthesis, except at Month 4 in which the controls accumulated significantly more sugars than the PGR treated plants (Figure 3a). Total sugars accumulated generally decreased significantly at later months until it reached a negative plateau at Month 5 to 6 (Figure 3a).

**Figure 3.**
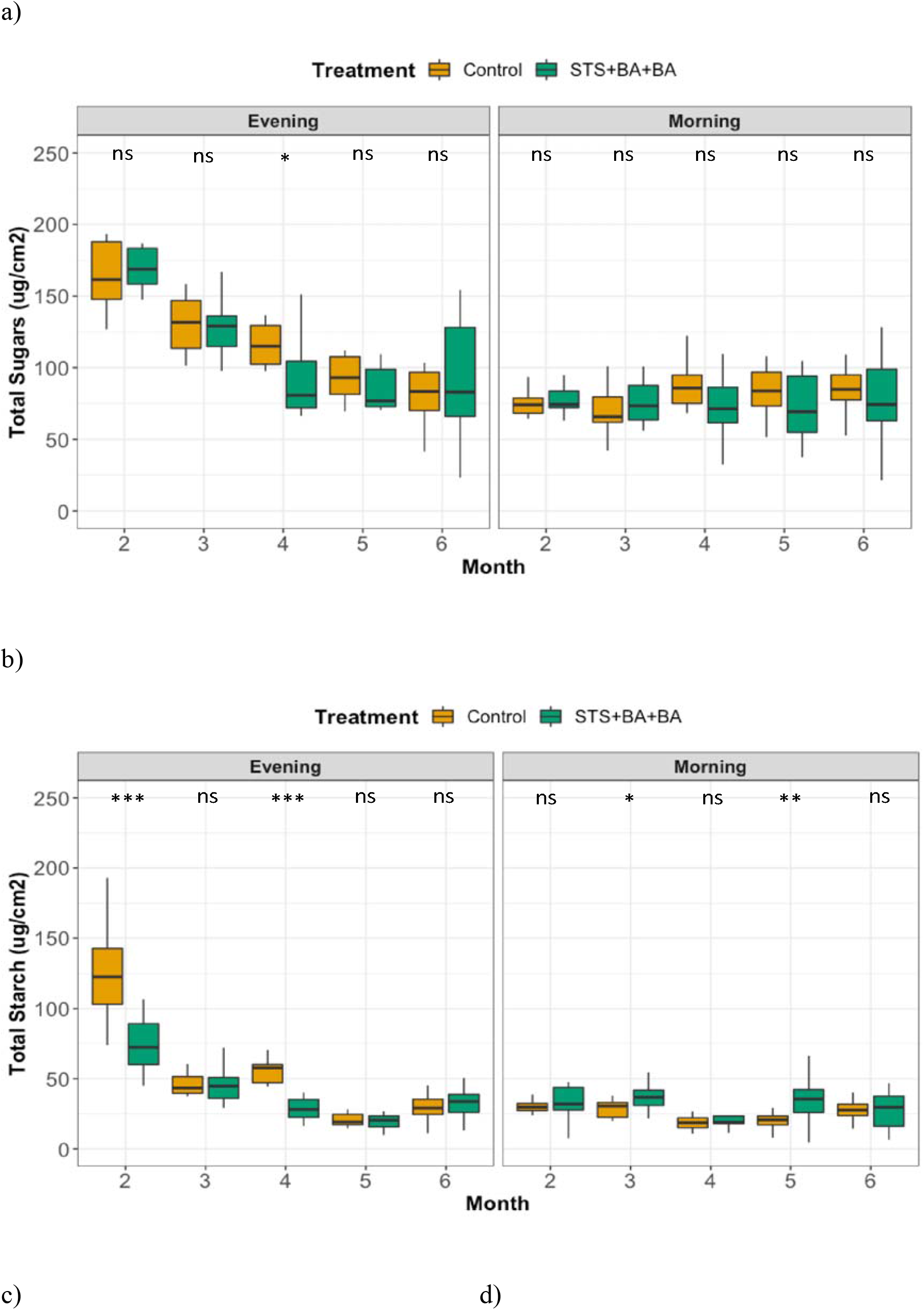
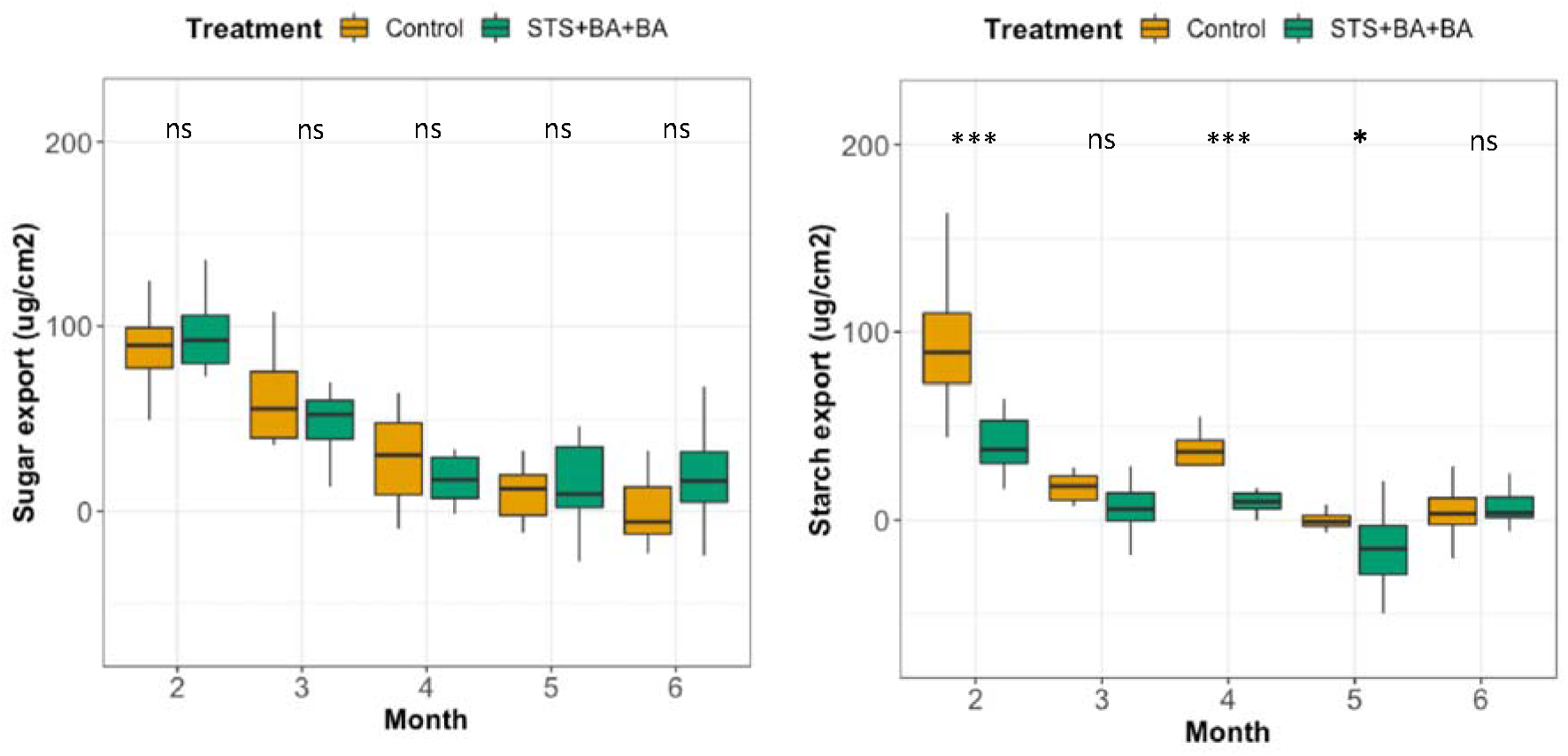
Cassava leaf carbohydrate accumulation, retention and export over time in PGR treated vs Control. a) Total sugars accumulated at evening; b) Total sugars retained the following morning (∼16 hrs later); c) Starch accumulated at evening; d) starch retained the following morning (∼16 hrs later); e)Total sugars exported (evening sugar – morning sugars); f) Starch exported (evening starch – morning starch). *, **, *** and ns indicate statistical significance on pairwise comparisons between treatments at 0.05, 0.01,and 0.001 significance levels, respectively.

Starch accumulation in recently matured leaves followed a general pattern similar to sugars (Figure 3c). At Month 2, prior to the start of storage root growth, starch accumulated during the day to high contents, then as storage root growth increased, the amount of starch accumulated was much less at later stages. Morning starch levels were low and stable throughout the time frame (Figure 3d). PGR treatment decreased the extent of day-time starch accumulation in leaves: At Months 2 and 4, evening starch levels were significantly lower in the PGR treatment than the control, while morning starch levels were significantly higher at Months 3 and 5 in the PGR treatment than the control (Figure 3d).

The difference between levels of leaf storage carbohydrates at the end of day and in the early morning was interpreted as largely representing phloem export, and hence is an indicator of plant source-sink status. Due to the greater amount of day-time leaf starch in the first few months than at later time frames, nighttime export was highest at Month 2 and decreased substantially to a plateau between months four and six (Figure 3e, f). While the amount of sugar exported was not significantly different between PGR-treated plants and controls (Figure 3e), starch export was significantly higher in controls than PGR-treated plants at months two, four and five (Figure 3f). It is notable that the evening and morning trends in starch levels that contributed to the lower amount of apparent export in PGR treated plants involved both a lesser amount of starch accumulation at the end of the day (Figure 3a), and a larger amount of starch remaining in the morning (Figure 3b). This suggests that PGR treatment possibly affects both photosynthesis in the daytime and sink demand at night.

### 3.4 Relationship between reproductive and vegetative development in cassava

During these studies it was apparent that many of the plants within a plot did not survive to the reproductive stage and did not produce marketable storage roots. To document this, we determined whether there was a significant difference in the number of plants that survived until annual harvest between treatments. About 70% of control plants survived while about 50% of PGR-treated plants survived (Figure 4 a). These findings indicate that while the PGR treatment had a pronounced favorable effect on reproductive development (Fig 1b-d), its associated phytotoxicity and leaf removal for petiole feeding apparently weakened the plants such that many plants succumbed to diseases and other stressful conditions (periodic flooding from heavy rains, onset of the dry season in Month 5) that were present in the field. Of the plants that survived, about 95% and 75% had storage roots in the controls and PGR-treated plants, respectively (Figure 4b).

**Figure 4.**
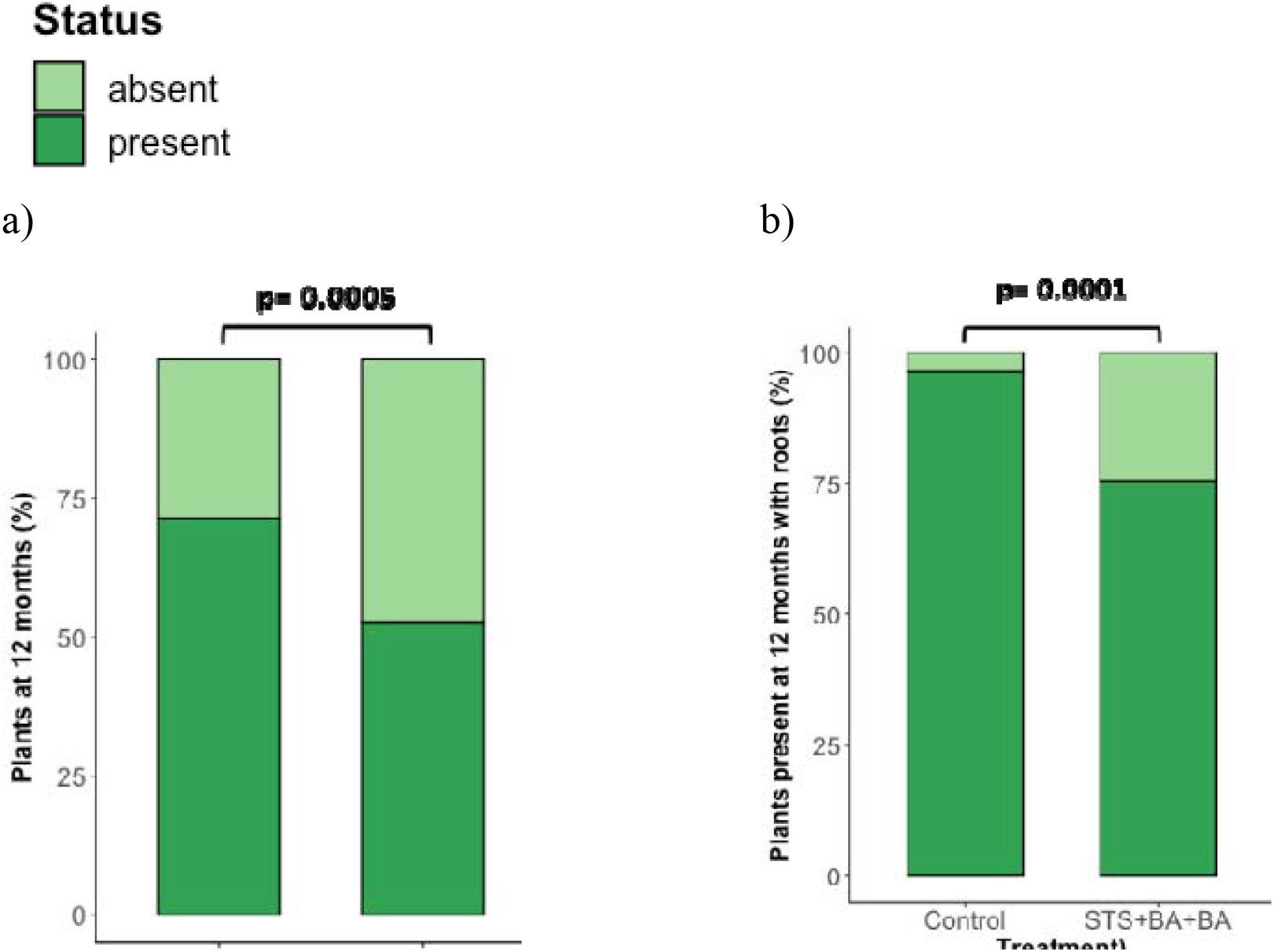
Percentage of plants (a) surviving at annual harvest, (b) survived and with roots P values reflect binomial model pairwise comparison of control versus PGR treatments.

## 4 Discussion

The current study showed that PGR treatment with anti-ethylene (STS) and cytokinin (BA) PGRs stimulated reproductive development such that the number of female flowers was increased by more than five-fold, and the number of fruits was increased three-fold. This is consistent with previous studies of cassava which have shown that STS stimulates more prolific inflorescence and flower development and extends flower longevity (Hyde et al. 2020). The current study also showed that STS + BA PGRs increased the proportion of flowers that were female, such that more than 90% were female. This is consistent with previous studies which have shown that cytokinin application increases the proportion of flowers that are female in *Jatropha curcas, Plukenetia volubilis*, and *Sapium sebiferum*, which like cassava are in the family Euphorbiaceae (Pan et al. 2014; Chen et al. 2014; Gangwar et al. 2018; Luo et al. 2020; Ni et al. 2018; Fröschle et al. 2017). On the other hand, STS + BA PGR treatment did not accelerate flowering time (Fig. 1a). Cytokinin has been reported to initiate flowering in several plant systems (Bonhomme et al. 2000; D’Aloia et al. 2011), though such effects have not been found in cassava. Our findings are also consistent with previous studies of STS application to cassava, which indicated STS does not affect the timing of flower initiation (Hyde et al. 2020). In contrast to the studies reviewed here, there have been no previous reports about the effect of co-application of STS and a synthetic cytokinin (Benzyladenine) on flowering time, particularly in cassava. In the current study, we showed that a mixture of STS and BA, applied by petiole feeding combined with BA sprayed to the shoot apex, did not accelerate flowering time, but it improved downstream reproductive events synergistically – increasing the number of total flowers and the fraction of flowers that are female, which in turn increased the number of fruits (Figure 1).

A possible explanation for stimulatory effects of anti-ethylene and cytokinin on the quantity of flowers and fruits is that these PGRs might alter plant source-sink balance and resource partitioning. While the small number and size of cassava flowers and fruits infers that they do not represent large sinks compared to the primary sinks, storage roots and vegetative shoots, it is plausible that these large competing sinks could limit the availability of photosynthate to flowers and fruits at a critical time in their development. We hypothesized that PGR treatment would decrease the strength of these competing sinks and decrease phloem flux to them so that more photosynthate would be available for flowers and fruits. This follows from some evidence that hormones affect partitioning among competing sinks. Root feeding of cytokinin and its transport to shoots has been shown to increase the shoot to root ratio in *Urtica dioica* (Fetene and Beck 1993). In carrot, foliar spray of the anti-ethylene STS significantly increased shoot growth and did not affect root growth over untreated plants when roots were restricted (Thomas 1993). However, in the current study, treatment of above-ground plant parts with a combination of STS and BA did not affect shoot or root growth and did not affect the proportion of root weight relative to total plant weight (Partitioning index, PI) and dry matter content in storage roots between Months 2 and 5 (Figure 2) in which reproductive development was vigorous (Figure 1). At Month 6, storage root and shoot fresh weight were higher in the PGR treatment than the control; however, these increases occurred after the main flowering episodes were complete and were likely due to the activation of repressed branch buds by cytokinin in treatment and so more lateral branches contributing to shoot weight. In addition, the lateral branches developed provided more leaves for photosynthesis relative to untreated plants at month 6. Although growth data in Figure 2 b are on a fresh weight basis, dry matter content of storage roots at Month 6 was slightly higher in PGR treatments than in controls (P= 0.0371) substantiating that dry matter growth in roots and shoots was greater in the PGR treatment, contrary to the hypothesis that PGR reduces growth of sinks that compete with flowers and fruits. Based on these findings, we conclude that the PGR effects on flower and fruit numbers was not because the treatment altered whole plant partitioning between storage roots and above-ground shoots.

Starch metabolism has previously been shown to follow a diurnal rhythm, with accumulation derived from photosynthesis peaking before dusk and degradation accompanied by export to sinks occurring at night; thus arriving at the lowest levels after dawn (Smith et al. 2005). Impairment to this process such as reduced starch accumulation or export or incorrect timing of rhythm is associated with decreased growth (McClung and Gutiérrez 2010). To some extent, leaf tissues can also temporarily store sugars with a diurnal trend. In current study starch export declined significantly under PGR treatment at Months 2, 4 and 5 (relative to the control) while sugar export was held at similar levels as the control (Figure 3). This decline is, however, not reflected in vegetative growth of either the shoots or roots (Figure 2). Furthermore, the carbohydrate export was not correlated with reproductive growth as treated plants had a greater number of flowers and fruits than the controls (Figure 1), which is expected to place more demand on photosynthate. This shows that in cassava the effect of PGR on flower and fruit numbers was not correlated with effects on the extent to which photosynthate is used for export at night and also that the PGR effects on cassava flower development is not due to nighttime photosynthate export.

While the combination of STS and BA provided considerable benefit to flower and fruit production, it had some phytotoxicity effects on leaves. Such phytotoxicity has been previously reported for STS (Serek et al. 2012; Høyer 1998; Hyde et al. 2020) in cassava and other species and it is generally tolerated given that the benefit for flower and fruit production outweighs the negative effects. In the current field application, however, PGR treatment significantly decreased the number of plants that survived until harvest at about 12 months. Treated plants that survived also had a decreased likelihood of having storage roots from which dry matter (indicative of storage roots being consumable) was obtainable due to rot or complete lignification (Figure 4). Under PGR treatments, plants were more susceptible to disease which negatively impacted growth later. Other studies have found that in some cases the mechanism of disease infection requires a destabilization of hormone balance for infection to proceed (Huang et al. 2018). We speculate this possibility occurred with respect treatments in current study. Given that the intended use of the PGR treatments is to assist breeding programs obtain better flower, fruit and seed production at first or second flowering tiers in the early part of the season, the plant mortality would seldom be a problem. Nevertheless, the current findings underscore that phytotoxicity is a potential problem, especially if applications extend for a relatively long timeframe.

## 5 Conclusion

This present study investigated the effect of a combination of STS and BA on cassava’s vegetative and reproductive growth to test the hypothesis that the benefit of STS + BA PGR treatments in increasing flower and fruit production are due, in part, to altered resource partitioning and decreased competition from storage roots and vegetative shoot sinks. However, PGR treatment did not decrease growth of storage roots or shoot vegetative growth. This indicates that the PGRs have effects more directly on flower and fruit reproductive signaling and regulatory pathways rather than an indirect effect on resource partitioning. We conclude that the PGR improvement of cassava flower and fruit development was not mediated through suppression of root and shoot sinks that compete for available photosynthate.

## 7 Author Contributions

D.O and T.S obtained funding. D.O and T.S designed experiment. D.O conducted field experiments while T.S conducted laboratory assays. D.O analysed data. D.O and T.S wrote manuscript.

## 9 Acknowledgments

The authors would also like to thank the Federal Government of Nigeria for partly funding Deborah’s studies through the Presidential Special Scholarship for Innovation and Development (PRESSID) managed by the National Universities Commission (NUC) and funded by the Federal Scholarship Board (FSB). The authors would also like to thank the Bioscience unit and Cassava Breeding unit of the International Institute of Tropical Agriculture, Nigeria for providing office space and field space for conducting experiments. The authors thank Olayemisi Esan, Peter Hyde and Peter Kulakow for useful conversations about this study.

This research was part of the Ph.D. dissertation of the first author at Cornell University, Ithaca, NY

