## Supplementary figures for "Enhancing cassava reproductive development does not negatively impact shoot to root ratio and dry matter content in storage roots"

### Supplementary data

a)

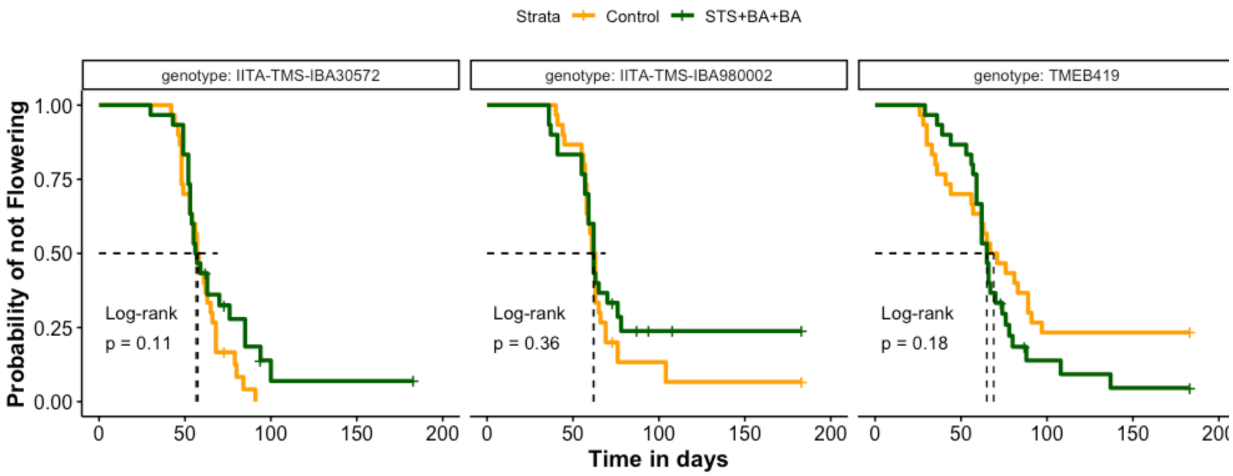


b)

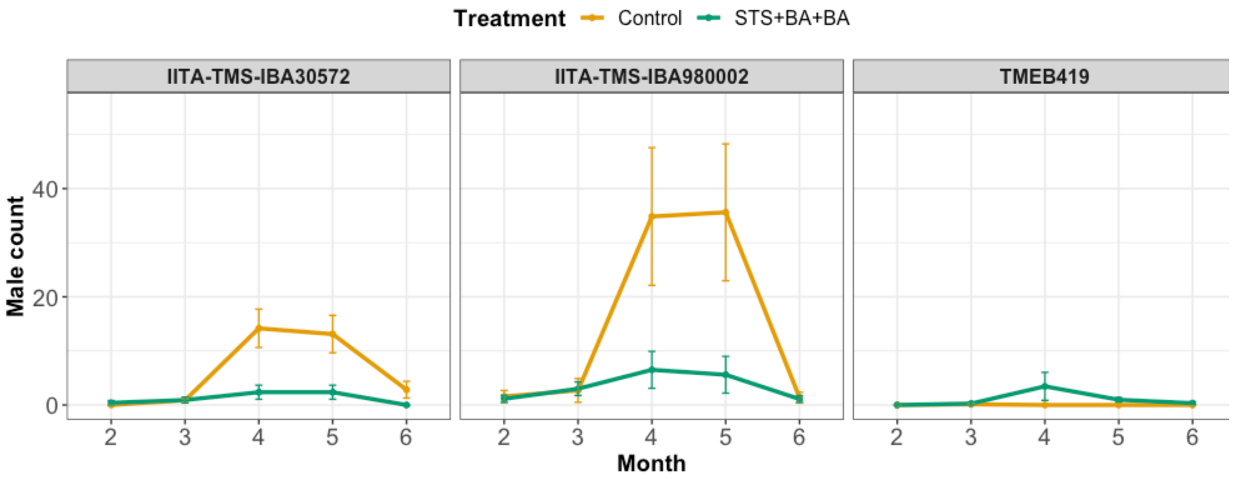


c)

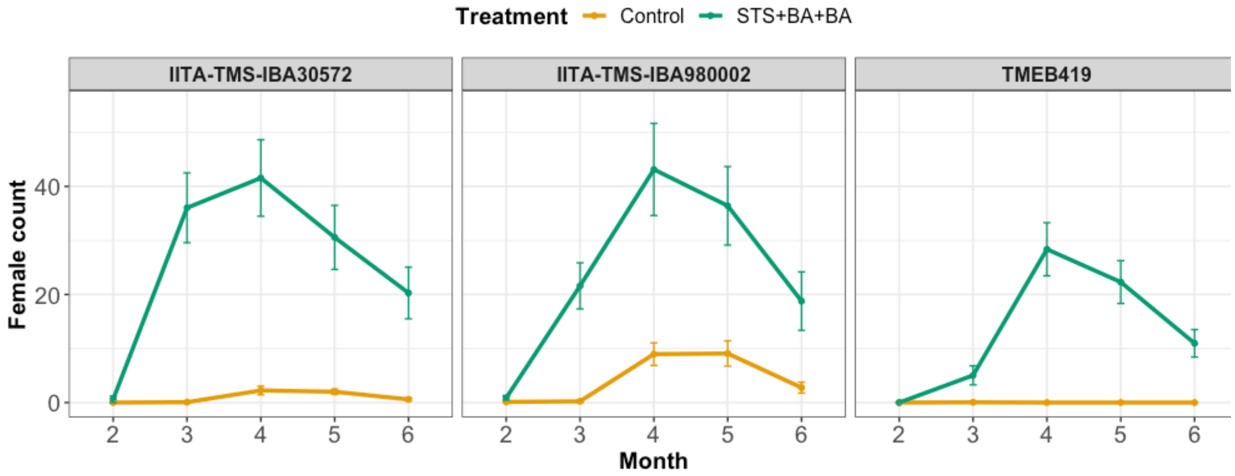


d)

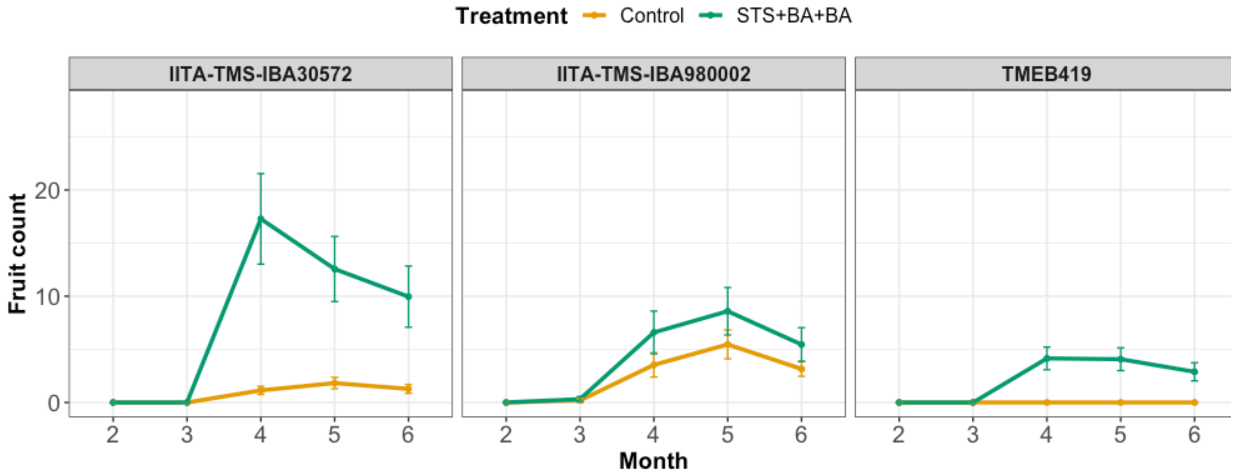


**S1** Genotypic responses of cassava reproductive development over time in PGR treated vs Control. a) Likelihood of flowering, Kaplan-Meier curves for flowering times. b) Maximum male flowers c) Maximum female flowers. d) Maximum fruit counted per month.

a)

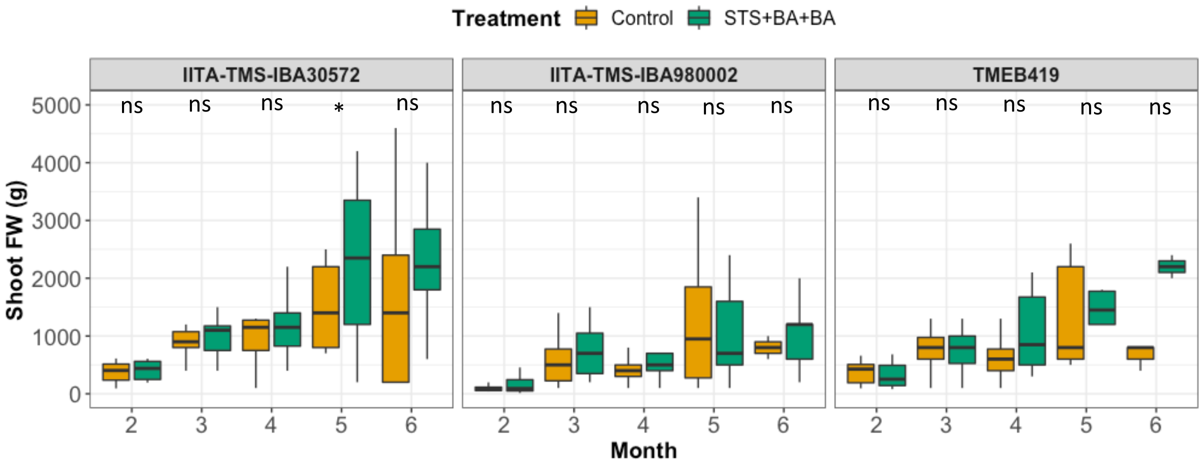

Treatment Control STS+BA+BA
IITA-TMS-IBA30572
IITA-TMS-IBA980002
TMEB419
Shoot FW (g)
Month
ns ns ns * ns
ns ns ns ns ns
ns ns ns ns ns


b)

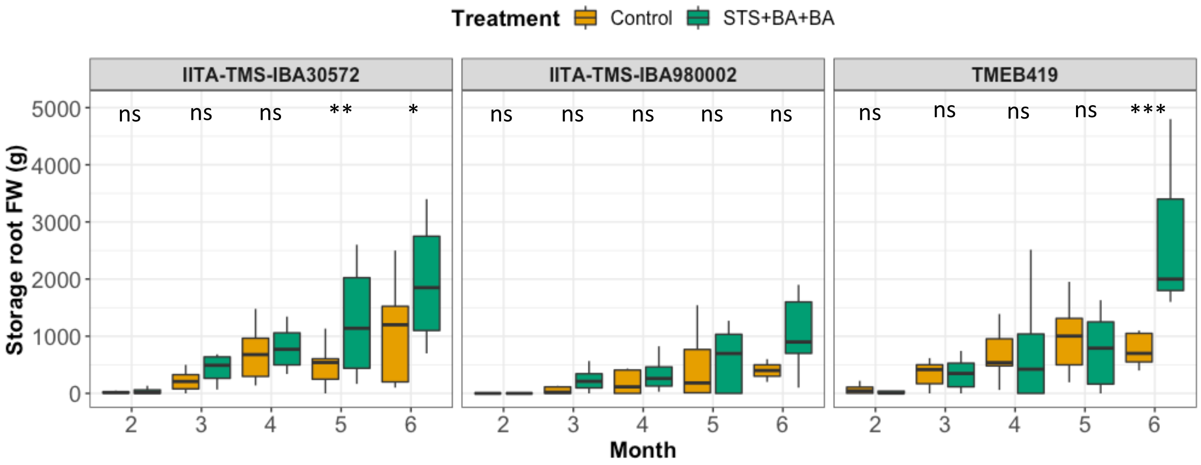

Treatment Control STS+BA+BA
IITA-TMS-IBA30572
IITA-TMS-IBA980002
TMEB419
Storage root FW (g)
Month
ns ns ns ** *
ns ns ns ns ns
ns ns ns ns ***


c)

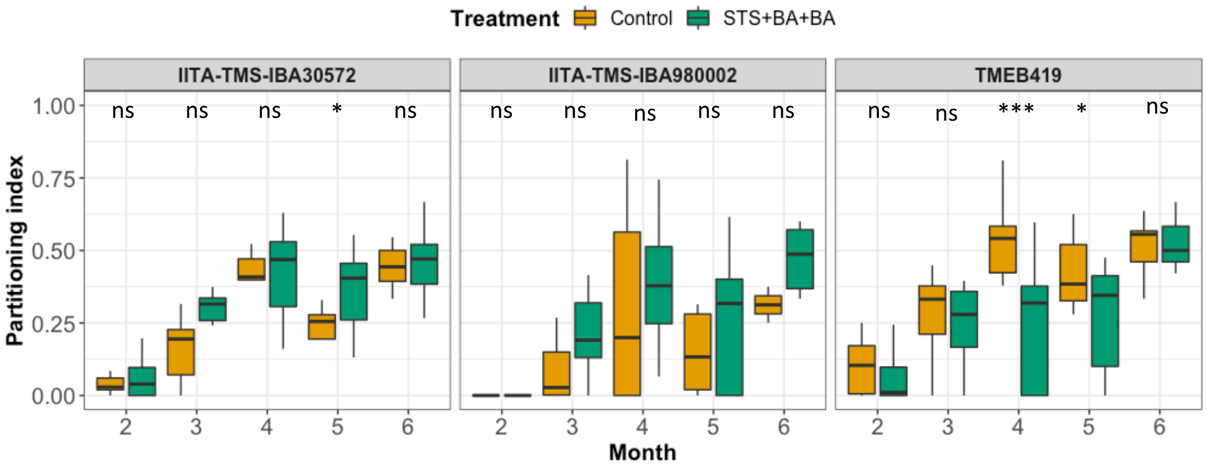

Treatment Control STS+BA+BA
IITA-TMS-IBA30572
IITA-TMS-IBA980002
TMEB419
Partitioning index
Month
ns ns ns * ns
ns ns ns ns ns
ns ns *** * ns

**d)**

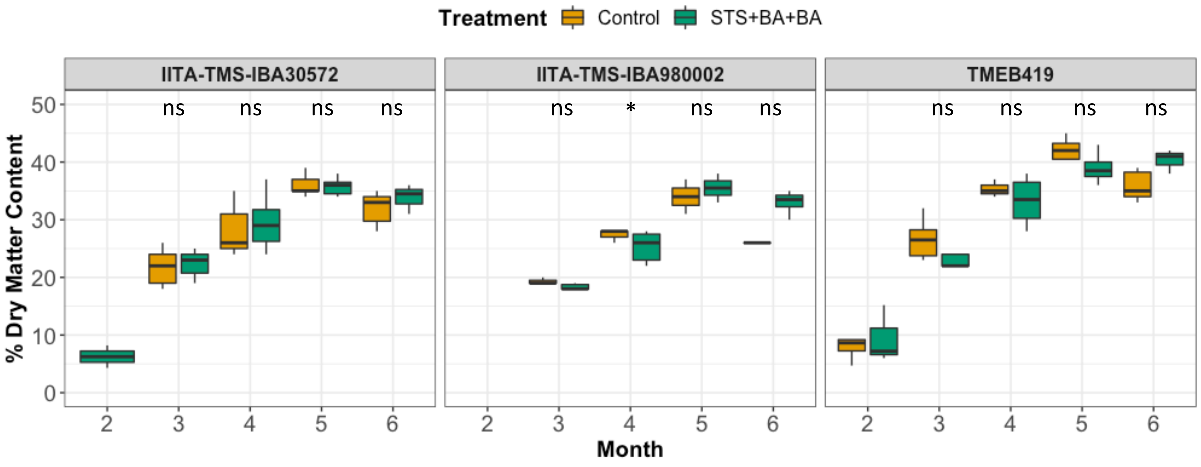


S 2 Genotypic responses of Vegetative growth over time over time in PGR treated vs Control. a) Shoot fresh weight b) Storage root fresh weight c) Partitioning index (on a fresh weight basis, see text) d) Percent dry matter content. *, and ns indicate statistical significance or no statistical significance, respectively, on pairwise comparisons between treatments (P<0.05).

**a)**

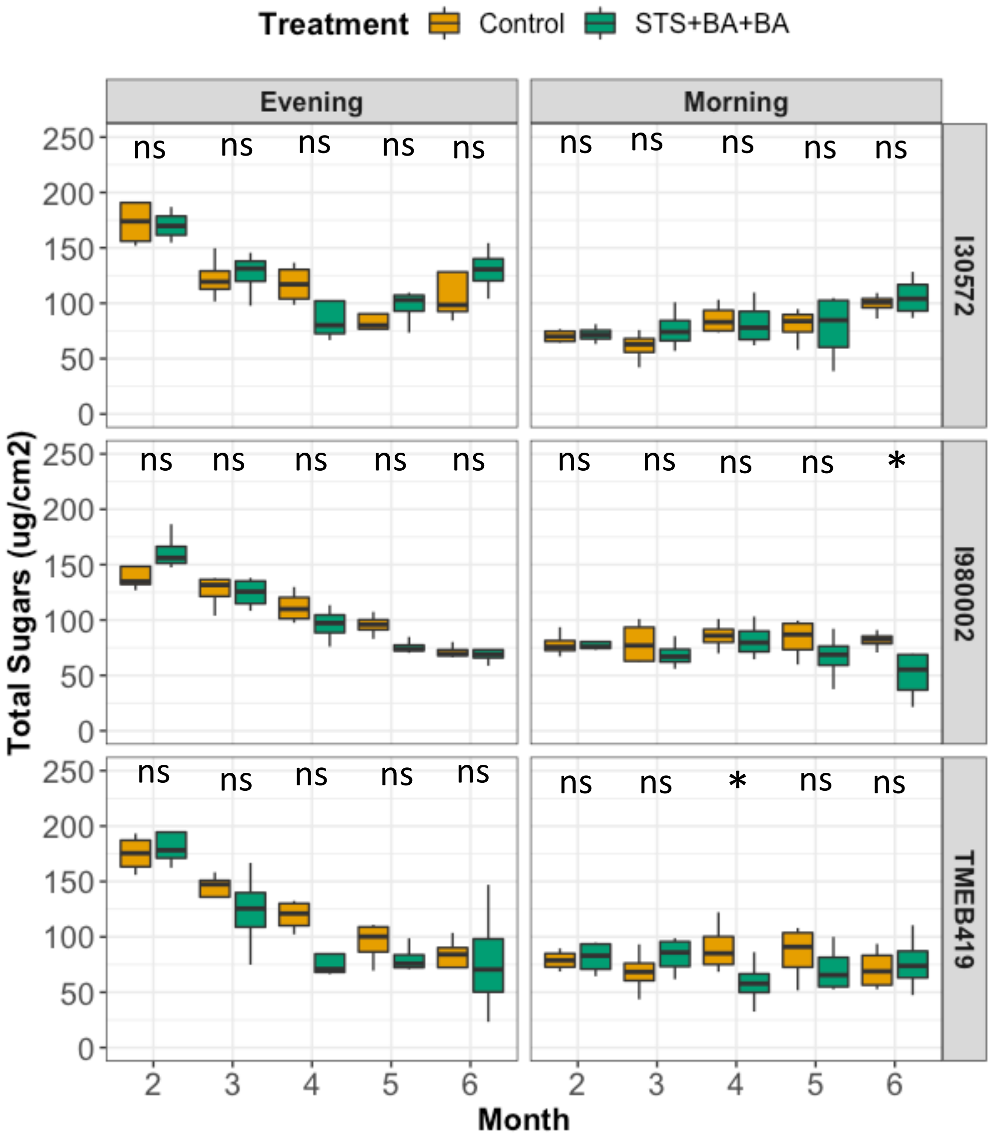

Treatment
Control
STS+BA+BA
Evening
Morning
I30572
I980002
TMEB419
Total Sugars (ug/cm2)
Month
ns
*

**b)**

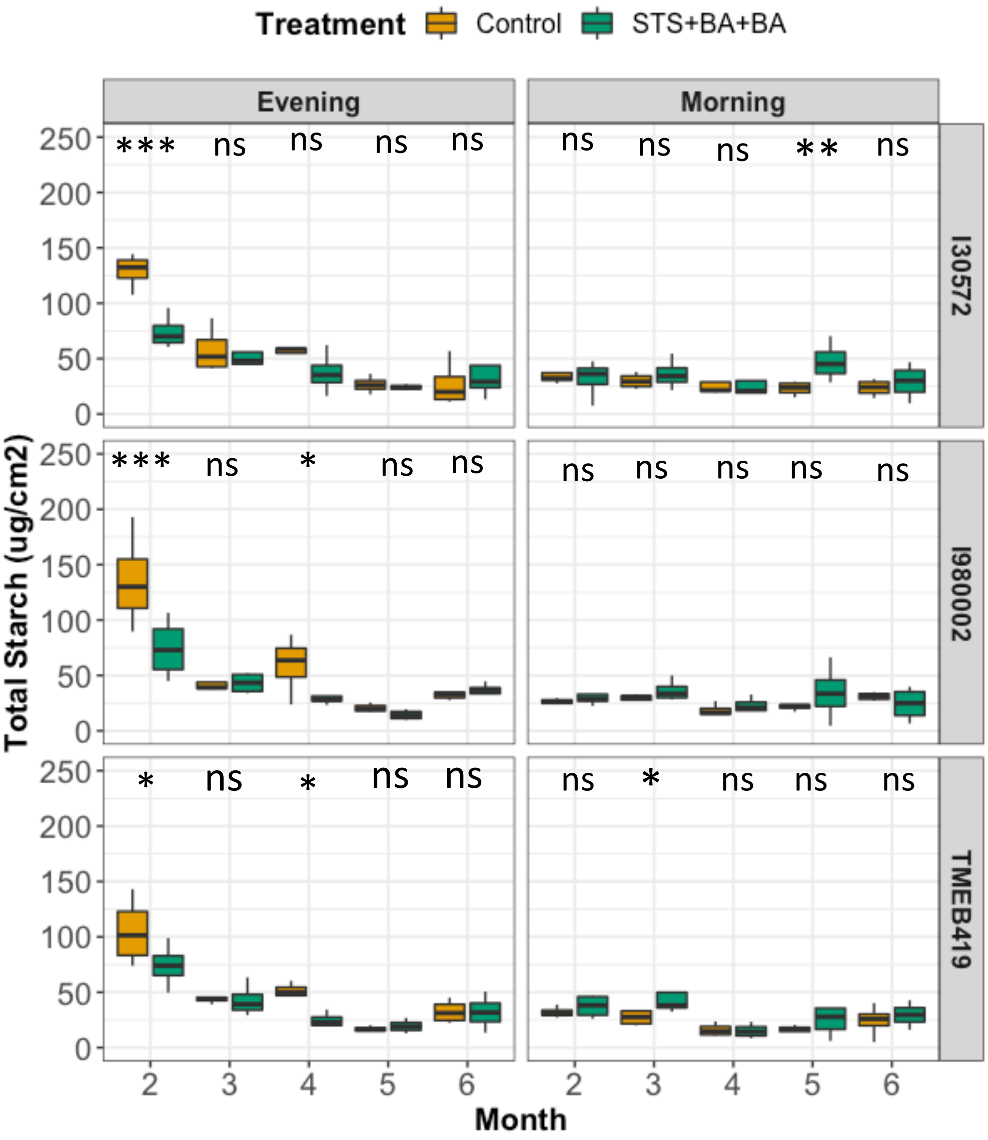
Treatment
Control
STS+BA+BA
Evening
Morning
I30572
I980002
TMEB419
Total Starch (ug/cm2)
Month
0
50
100
150
200
250
2
3
4
5
6
***
ns
**
*

c)

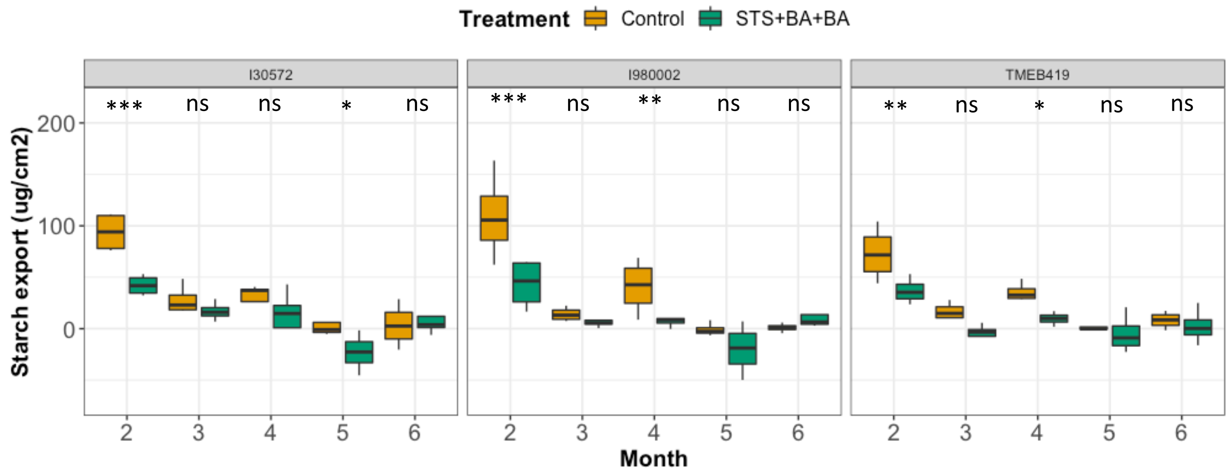


d)

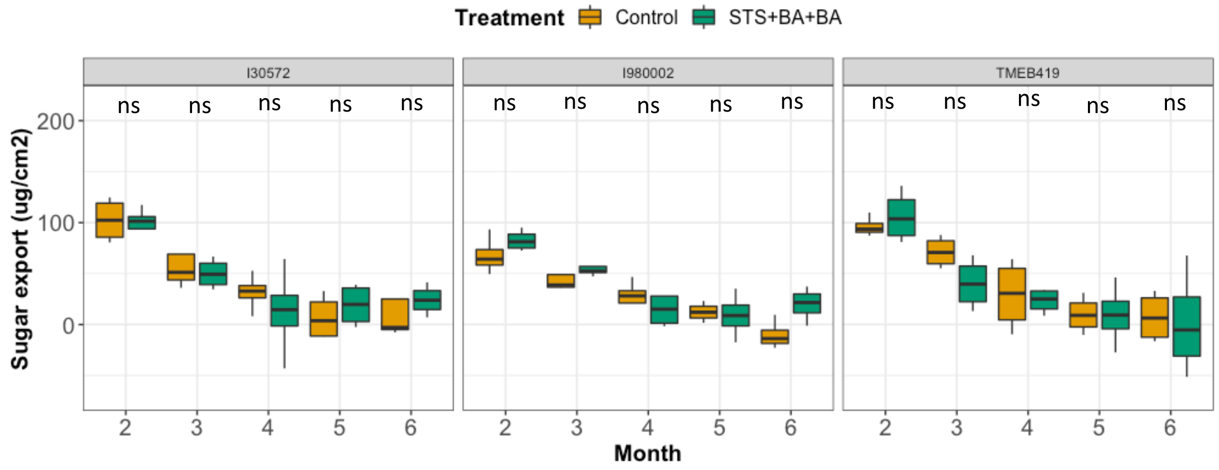


**S 3** Genotypic responses of cassava leaf carbohydrate accumulation, retention and export over time in PGR treated vs Control. a) Total sugars accumulated at evening; b) Total sugars retained the following morning (~16 hrs later); c) Starch accumulated at evening; d) starch retained the following morning (~16 hrs later); e)Total sugars exported (evening sugar – morning sugars); f) Starch exported (evening starch – morning starch). *, ** ,*** and ns indicate statistical significance on pairwise comparisons between treatments at 0.05, 0.01,and 0.001 significance levels, respectively.

a)

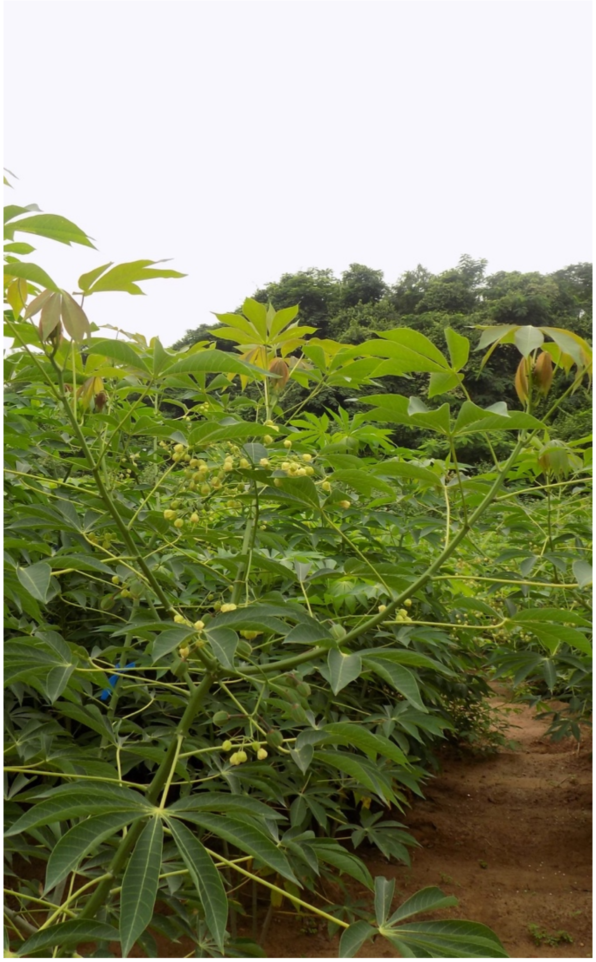

b)

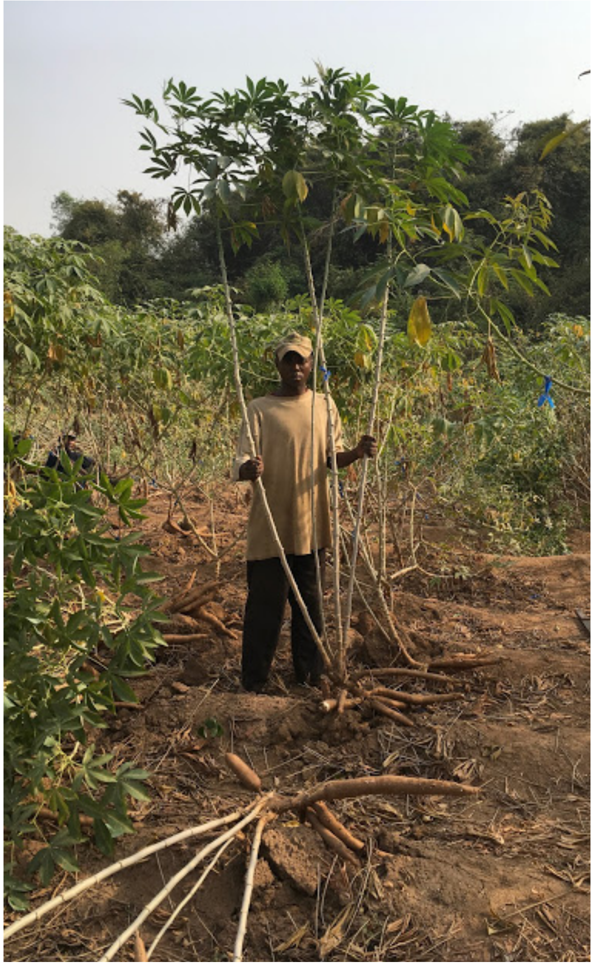

**S 4** Cassava grown on the field a) at 3 months after planting ('0002) b) at 7 months after planting ('419)
